# Impaired phloem loading in genome-edited triple *knock-out* mutants of SWEET13 sucrose transporters

**DOI:** 10.1101/197921

**Authors:** Margaret Bezrutczyk, Thomas Hartwig, Marc Horshman, Si Nian Char, Jinliang Yang, Bing Yang, Wolf B. Frommer, Davide Sosso

**Affiliations:** Institute for Molecular Physiology, Heinrich Heine University Düsseldorf, and Max Planck Institute for Plant Breeding Research, Cologne, Germany; Department of Plant Biology, Carnegie Science, 260 Panama St., Stanford, CA 94305, USA; Department of Genetics, Development, and Cell Biology, Iowa State University, Ames, IA 50011, USA; Department of Agronomy and Horticulture, University of Nebraska-Lincoln, Lincoln, NE 68588, USA

**Keywords:** corn, efflux, export, leaf, phloem loading, maize, sucrose, transport

## Abstract

Crop yield depends on efficient allocation of sucrose from leaves to seeds. In *Arabidopsis*, phloem loading is mediated by a combination of SWEET sucrose effluxers and subsequent uptake by SUT1/SUC2 sucrose/H^+^ symporters. ZmSUT1 is essential for carbon allocation in maize, but the relative contribution to apoplasmic phloem loading and retrieval of sucrose leaking from the translocation path is not known. We therefor tested whether SWEETs are important for phloem loading in maize.

Here we identified three leaf-expressed *SWEET* sucrose transporters as key components of apoplasmic phloem loading in *Zea mays* L. Notably, *ZmSWEET13* paralogs (*a*, *b, c*) are among the highest expressed genes in the leaf vasculature. Genome-edited triple *knock-out* mutants are severely stunted. Photosynthesis of mutants was impaired and leaves accumulated starch and soluble sugars. RNA-seq revealed profound transcriptional deregulation of genes associated with the photosynthetic apparatus and carbohydrate metabolism. GWAS analyses may indicate that variability in *ZmSWEET13s* is correlated with agronomical traits, specifically flowering time and leaf angle.

This work provides support for cooperation of three ZmSWEET13s with ZmSUT1 in phloem loading in *Zea mays* L. Our study highlights these three ZmSWEET13 sucrose transporters as possible candidates for the engineering of crop yield.

**One Sentence Summary:** Three *SWEET* sucrose transporter paralogs are necessary for phloem loading in maize.

## Introduction

Crop yield is critical for human nutrition, yet the underlying machinery that ultimately determines yield potential is still not understood. Crop productivity under ideal conditions is determined by the efficiency with which plants intercept light, convert it into chemical energy, translocate photosynthates, and convert these to storage products in harvestable organs (Zhu *et al.*, 2010). In many crops, sucrose is the primary form for translocation inside the conduit (the phloem). A combination of SWEET-mediated efflux from phloem parenchyma and subsequent secondary active sucrose import by SUT sucrose/H^+^ symporters is thought to create the driving force for pressure gradient-driven phloem transport and retrieval of sucrose leaking along the translocation path (Chen *et al.*, 2015a).

There is a debate regarding the mechanisms of phloem loading in crops. Sucrose is thought to follow one of three routes: (i) apoplasmic loading via plasma membrane transporters, (ii) symplasmic loading via diffusion through plasmodesmata, or (iii) polymer trapping via enzymatic addition of galactinol, which is thought to impair back-diffusion through plasmodesmata (Turgeon & Wolf, 2009; Chen *et al.*, 2015a). Some mechanisms may coexist, as suggested by anatomical studies which have found thin and thick walled sieve tubes in monocots, cell types that may differ regarding the primary loading mechanism (Botha, 2013).

In *Arabidopsis*, a SWEET/SUT-mediated apoplasmic mechanism appears important for phloem loading (Chen *et al.*, 2012, 2015a). SWEETs are a class of seven transmembrane helix transporters that function as hexose or sucrose uniporters (Xuan *et al.*, 2013). Multiple SemiSWEETs and SWEETs have been crystallized, and AtSWEET13 has been proposed to function in complexes via a ‘revolving door’ mechanism to accelerate transport efficacy (Feng & Frommer, 2015; Latorraca *et al.*, 2017; Han *et al.*, 2017). In Arabidopsis, SWEET roles include phloem loading, nectar secretion, pollen nutrition, and seed filling (Chen *et al.*, 2012; Sun *et al.*, 2013; Lin *et al.*, 2014; Sosso *et al.*, 2015). In rice, cassava, and cotton, SWEETs act as susceptibility factors for pathogen infections (Chen *et al.*, 2010; Cohn *et al.*, 2014; Cox *et al.*, 2017). AtSWEET11 and 12 are likely responsible for effluxing sucrose from the phloem parenchyma into the apoplasm (Chen *et al.*, 2012). Sucrose is subsequently loaded against a concentration gradient into the SECC via the SUT1 sucrose/H+ symporter (a.k.a. AtSUC2), powered by the proton gradient created by co-localized H+/ATPases (Riesmeier *et al.*, 1994; Gottwald *et al.*, 2000; Slewinski *et al.*, 2009; Srivastava *et al.*, 2009). Though the fundamental involvement of SUT transporters in phloem loading has been demonstrated using RNAi and knock-out mutants in *Arabidopsis* (also in potato, tobacco, tomato, and maize (Riesmeier *et al.*, 1994; Bürkle *et al.*, 1998; Srivastava *et al.*, 2009; Chen *et al.*, 2015a), *atsweet11,12* and *atsuc2 (sut1)* mutants only showed partially impaired phloem loading.

In monocots, including all cereal crops, the situation is less clear. In maize, the phloem-expressed ZmSUT1 (Baker *et al.*, 2016) (phylogenetically in the SUT2 clade) appears to be critically important for phloem translocation (Slewinski *et al.*, 2009), whereas rice *ossut1* mutants and RNAi lines had no apparent growth or yield defects (Ishimaru *et al.*, 2001; Scofield *et al.*, 2002; Eom *et al.*, 2012). As a result, there is an ongoing debate regarding the mechanisms behind phloem loading in cereals (Braun *et al.*, 2014; Regmi *et al.*, 2016).

Here we identified a set of three close paralogs of SWEET13 from *Z. mays* L. as essential sucrose transporters for phloem loading.

## Material and Methods

### Plant material and growth conditions

*zmsweet13a*, *zmsweet13b* and *zmsweet13c* alleles were obtained with a CRISPR-Cas9 construct targeting a sequence (5’-GCATCTACAAGAGCAAGTCGA<u>CGG</u>-3’, the underlined CGG for PAM) conserved in all three paralogs in the 3rd exon as described (Char *et al.*, 2017). T0 plants were selfed or outcrossed to B73 and plants which did not contain the CRISPR construct were selected for further analysis. T1, T2, and T3 plants homozygous for all three mutated genes (*zmsweet13abc*) were selected along with wild-type siblings. Height was assessed by weekly measurement from the soil surface to the top of the highest fully-developed leaf. Wild-type “siblings” were descendants of the Hi-II plants transformed and outcrossed once to B73, which in the T1 generation did not carry the CRISPR-Cas9 construct or any detectable mutations. Triple mutant plants either descended from selfed T0 Hi-II plants or outcrossed once to B73. The mutant phenotype was unaffected by the genetic difference. Mutants and wild-type plants were grown side by side, in greenhouses under long-day conditions (16h day/8h night, 28-30 °C), and in 2016 in a summer field at Carnegie Science (Stanford, California, USA).

### Genotyping of rice and maize plants

Genomic DNA was extracted from leaves using a Qiagen Biosprint 96. PCR was performed with the Terra PCR Direct Red Dye Premix Protocol (Clontech Laboratories) with melting temperatures of 60 °C, 64 °C, and 62.5 °C for *ZmSWEET13a*, *b*, and *c,* respectively (for primers see Table S2). Amplicons of relevant regions of the CRISPR-Cas9 targeted *ZmSWEET13* alleles were sequenced by Sequetech (Mountain View, CA). Chromatograms were analyzed using 4Peaks (www.nucleobytes.com/4peaks/).

### Plastic embedding and sectioning

Flag leaves collected at 7:00 am were placed in 0.1 M cacodylate buffered fixative with 2% paraformaldehyde and 2% glutaraldehyde, vacuum infiltrated for 15 min and incubated overnight. Sample dehydration was performed by a graded ethanol series (10%, 30%, 50%, 70% and 95%). Sample embedding was performed according to the LR White embedding kit protocol (Electron Microscopy Science). Cross-sections (1.5 µm) were obtained on an Ultracut (Reichert), stained 30s with 0.1% toluidine blue and washed with ddH_2_O (2x), followed by 5 min of starch staining with saturated Lugol’s solution. Sections were mounted with CytoSeal 60 (EM Science).

### Phylogenetic analyses

The evolutionary history was inferred by using Maximum Likelihood with a JTT matrix-based model. The tree with the highest log likelihood (-3000.1) is shown. The percentage of trees in which associated taxa clustered together is shown next to the branches. Initial tree(s) for the heuristic search were obtained by Neighbor-Joining to a matrix of pairwise distances with the JTT model used for estimation. The analysis involved 16 polypeptides sequences. A minimum of 95% site coverage was required so that no more than 5% alignment gaps, missing data, and ambiguous bases were allowed at any position. There were a total of 252 positions in the final dataset. Evolutionary analyses were conducted in MEGA6.

### Soluble sugar analyses

Flag leaves were harvested from mature plants at 7:00 am. 70 mg of liquid nitrogen-ground tissue was incubated for 1 hour with 1 ml of 80% ethanol on ice with frequent mixing. Samples were spun for 5 min at 4 °C at 13,000 *g*, and supernatant was removed. This step was repeated once. The liquid supernatant was subsequently dried in a vacuum concentrator and re-suspended in water. Sucrose, glucose and fructose were measured using NAD(P)H-coupled enzymatic methods using a plate reader M1000 (Tecan), with measured values normalized to fresh weight. Starch quantification was performed as previously described (Sosso *et al.*, 2015).

### Starch staining

Flag leaves collected at 7:00 am were were boiled in 95% ethanol for approximately 30 min (until chlorophyll pigments disappeared). Cleared leaves were submerged in saturated Logol’s iodine (IKI) solution for 15 min, rinsed twice with H_2_O, and imaged with a Lumix GF1 camera (Panasonci, Kadoma, Osaka, Japan). The IKI solution used for starch staining was made by adding 1 g of iodine and 1 g of Potassium Iodide to 100 mL H_2_O.

### RNA isolation and transcript analyses

RNA was extracted using the Trizol method (Invitrogen). First strand cDNA was synthesized using Quantitect reverse transcription Kit (Qiagen). qRT-PCR to determine expression level was performed using the LightCycler 480 (Roche), and the 2^-ΔCt^ method for relative quantification. Wild-type maize and *zmsweet13abc* flag leaves were sampled at 5:00 pm. Primers in the last exon and the 3′ UTR of *ZmSWEET13a, b,* and *c* (Supplementary Table 2) were used for qRT-PCR to determine gene expression levels. Internal references were *Zm18s* and *ZmLUG*.

### FRET sucrose sensor analysis in HEK293T cells

*ZmSWEET13a, b,* and *c* coding sequences were cloned into the Gateway entry vector pDONR221f1, followed by LR recombination into pcDNA3.2V5 for expression in HEK293T cells. HEK293T cells were co-transfected with *ZmSWEET13a, b,* or *c* in *pcDNA3.2V5* and the sucrose sensor FLIPsuc90µΔ1V (Chen *et al.*, 2012) using Lipofectamine 2000 (Invitrogen). For FRET imaging, HBSS medium was used to perfuse HEK293T/FLIPsuc90µΔ1V cells with defined pulses containing 20 mM sucrose in buffer. Image acquisition and analysis were performed as previously described (Chen *et al.*, 2012). AtSWEET12 was used as a positive control. Negative control were empty vector transfectants.

### Transient gene expression in *Nicotiana benthamiana* leaves

The *Agrobacterium tumefaciens* strain GV3101 was transformed with the binary expression clone (pAB117) carrying *ZmSWEET13a*, *b*, or *c* C-terminally fused with eGFP and driven by the CaMV 35S promoter. Agrobacterium culture and tobacco leaf infiltration were performed as described (Sosso *et al.*, 2015). Chloroplast autofluorescence was detected on a Leica TCS SP8 confocal microscope (470 nm excitation with simultaneous 522–572 nm (eGFP) and 667–773 nm emission (autofluorescence)). Epidermal leaf chloroplast autofluorescence (Dupree *et al.*, 1991) allowed us to determine eGFP vacuolar localization (surrounding the chloroplasts) and plasma membrane localization was deduced (adjacent to chloroplasts; according to bright-field image). Image analysis done using Fiji software (https://fiji.sc/).

### Analyses of photosynthetic rates

Licor LI-6800 measurements were taken at mid-day under normal greenhouse conditions (28 °C, PAR 1000, 60% relative humidity). 2 cm diameter disc of leaf was clamped inside the Licor measurement chamber and relative levels of CO2 inside and outside of the chamber were measured, with µmol m^−2^s^−1^ CO_2_ absorbed by leaf segment inside chamber used as a proxy for photosynthesis rate. Measurements were made at the tips of Leaf 7 - Leaf 10 at mid-day.

### Candidate gene association study

To test whether sequences at SWEET loci are associated with phenotypic variations in the maize population, we analyzed the maize diversity panel composed of 282 inbred lines (HapMap3 SNP data (Bukowski *et al.*, 2015) for the panel from the Panzea database (www.panzea.org)). We filtered SNP data (MAF > 0.1; missing rate < 0.5) using PLINK (Purcell *et al.*, 2007) and calculated a kinship matrix using GEMMA (Zhou & Stephens, 2012) using the filtered SNP set. GWAS was performed by fitting a mixed linear model using GEMMA, where the kinship matrix was fitted as random effects in the model. An FDR approach (Benjamini & Hochberg, 1995) was employed to control the multiple test problem with a cutoff of 0.05. Linkage disequilibrium of SNPs in our candidate genes with significant association SNPs was calculated using PLINK (Purcell *et al.*, 2007).

### RNA-seq and data analysis

*zmsweet13abc* triple mutants and WT siblings were grown in soil under greenhouse conditions. Total RNA was isolated from flag leaf tissues using acidic phenol extraction as described previously (Eggermont *et al.*, 1996). Purification of poly-adenylated mRNA using oligo(dT) beads, construction of barcoded libraries, and sequencing using Illumina HiSeq technology (150 bp paired-end reads) were performed by Novogene (https://en.novogene.com/) using manufacturer recommendations. Trimmed and QC-filtered sequence reads were mapped to the B73 AGPv3 genome using STAR (v. 2.54) (Dobin *et al.*, 2013) in two pass mode (parameters: --outFilterScoreMinOverLread 0.3, -outFilterMatchNminOverLread 0.3, --outSAMstrandField intronMotif, --outFilterType BySJout, --outFilterIntronMotifs RemoveNoncanonical, --quantMode TranscriptomeSAM GeneCounts). Unique reads were filtered by mapping quality (q20) and PCR duplicates removed using Samtools (v. 1.3.1). Gene expression was analyzed in R (v. 3.4.1) using DEseq2 software (v. 1.16.1) (Love *et al.*, 2014). Genes were defined as differentially expressed by a two-fold expression difference with a p-value, adjusted for multiple testing, of < 0.05. Accession numbers for the RNA-Seq data in the Gene Expression Omnibus database will be made available.

## Results

To directly test whether SWEETs are involved in phloem loading in maize, we evaluated the role of leaf-expressed maize *SWEETs* in carbon allocation. We identified three *SWEET13* paralogs (GRMZM2G173669: *ZmSWEET13a*, GRMZM2G021706: *ZmSWEET13b*, GRMZM2G179349: *ZmSWEET13c*) as the most highly expressed SWEETs in maize leaves (**Fig. S1**). *ZmSWEET13a* and *b* are located in tandem on chromosome 10 in a region syntenic with the *OsSWEET13* locus in rice, while *ZmSWEET13c* is on chromosome 3 (**Fig. S2**). Interestingly, maize appeared to be one of few cereals carrying three *SWEET13* paralogs in its genome, along with *S. bicolor* and *T. urartu* (**Figure 1a; Fig. S3**). *ZmSWEET13a, b* and *c* are preferentially expressed in bundle sheath/vein preparations rather than mesophyll (**Fig. S4**), as is *ZmSUT1*. ZmSUT1 and the three SWEETs showed higher expression in leaf tips (**Fig. S5**). We tested the transport activity of the three SWEETs by coexpressing each with sucrose FRET (Förster resonance energy transfer) sensors in human HEK293T cells (Chen *et al.*, 2010, 2012). All three SWEETs mediated sucrose transport in mammalian cells (**Fig. 1b**). To test whether these SWEETs were part of (i) intercellular translocation or (ii) intracellular sugar sequestration similar to the *Arabidopsis* SWEET2, 16 or 17 (Chardon *et al.*, 2013; Klemens *et al.*, 2013; Guo *et al.*, 2014; Chen *et al.*, 2015b), we tested their subcellular localization in transiently transformed tobacco cells, and found that they localized preferentially to the plasma membrane (**Fig. 1c**).

**Figure 1.**
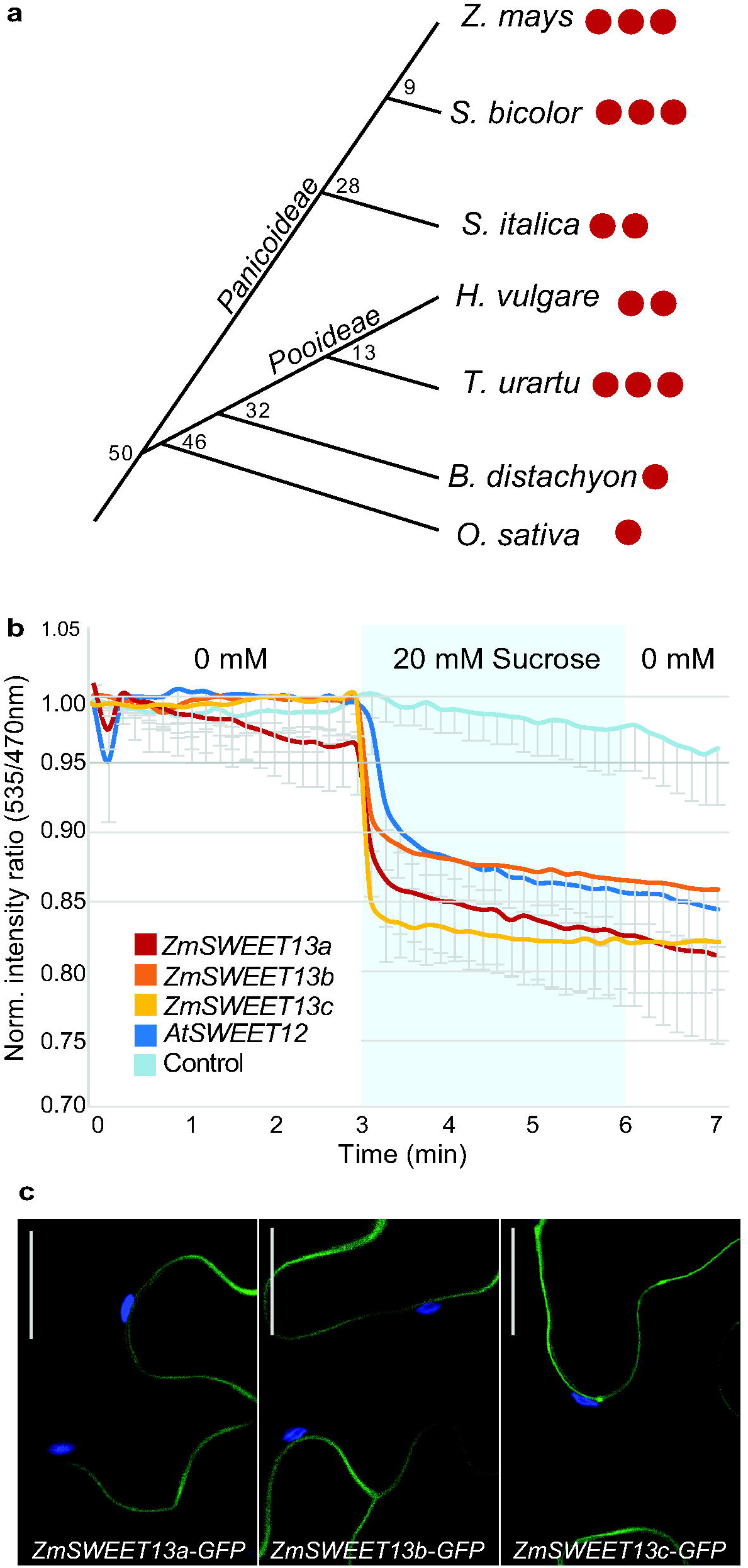
*ZmSWEET13* paralogs phylogeny, functional and subcellular characterization. **(A)** Scheme of phylogenetic relationship between putative orthologs in grasses. Chronogram branch divergence time-points are in million years (Emms *et al.*, 2016). **(B)** Sucrose transport activity by *ZmSWEET13a*, *b* and *c* in HEK293T cells coexpressing the FLIPsuc90µΔ1V (sucrose). Cell were transfected to express sensors only as negative control, or to co-express AtSWEET12 as positive controls for sucrose. HEK cells were subjected to a 20mM sucrose pulse for 3 minutes, (mean ± s.e.m., repeated independently four times with comparable results). **(C)** Confocal images (Z-stack) of *Agrobacterium*-infiltrated *N. benthamiana* epidermal leaf cells. ZmSWEET13a-eGFP (C-term), as well b and c indicated localization at the plasma membrane. The eGFP signal (Green; 522–572 nm) was merged with fluorescence derived from chloroplasts (Blue; 667–773 nm). Fluorescence was visualized using confocal laser scanning microscopy, 3 days after *Agrobacterium* infiltration. Scale bars, 50 µm.

Recently, *ZmSWEET13* had been implicated as a possible key player in C4-photosynthesis in grasses (Emms *et al.*, 2016). To test their role in maize, we designed guide RNAs that target a conserved region within a transmembrane domain, assuming that such mutations would lead to complete loss of function. We generated single *knock-out* mutants, as well as combinations of mutant alleles, using CRISPR-Cas9 (**Fig. S6**). Genome editing allowed us to recover two mutant alleles of *ZmSWEET13a*, four of *ZmSWEET13b* and three for *ZmSWEET13c*. The majority of mutations were caused by single nucleotide insertions in the target sequence. All mutations create premature STOP codons leading to truncated polypeptides (at amino acid 129 in the fourth of seven transmembrane domains) (**Fig. S6**). T2 lines carrying homozygous mutations in all three genes were characterized by severe growth defects (**Fig. 2a**). The growth phenotype was verified in subsequent generations in the greenhouse and a single field season. Single and double mutants showed slight growth defects, while triple mutants showed substantial defects: plants were severely stunted with shorter, narrower leaves (**Fig. 2a-c**). Leaves were chlorotic, accumulated 5x more starch and 4x more soluble sugars compared to wild-type (**Fig. 3a-c**), likely a consequence of impaired phloem loading. Accumulation of sugars and starch occurred primarily in mesophyll and bundle sheath cells and strongly impacted photosynthesis, even in greenhouse-grown mutant plants (**Fig. 3d, e; Fig. S7**). In the field, triple mutants from five independent allelic combinations presented even more severe phenotypes, with extreme chlorosis, massive anthocyanin accumulation and extremely stunted growth; in several cases resulting in lethality (**Fig. S8**). *SWEET13* mRNA levels were drastically reduced in all three *ZmSWEET13s*, as quantified by RNA-seq and qRT-PCR (**Fig. S9** and **Fig. 2d**). In summary, the strong phenotype of the triple mutant is consistent with maize using predominantly an apoplasmic phloem loading mechanism.

**Figure 2.**
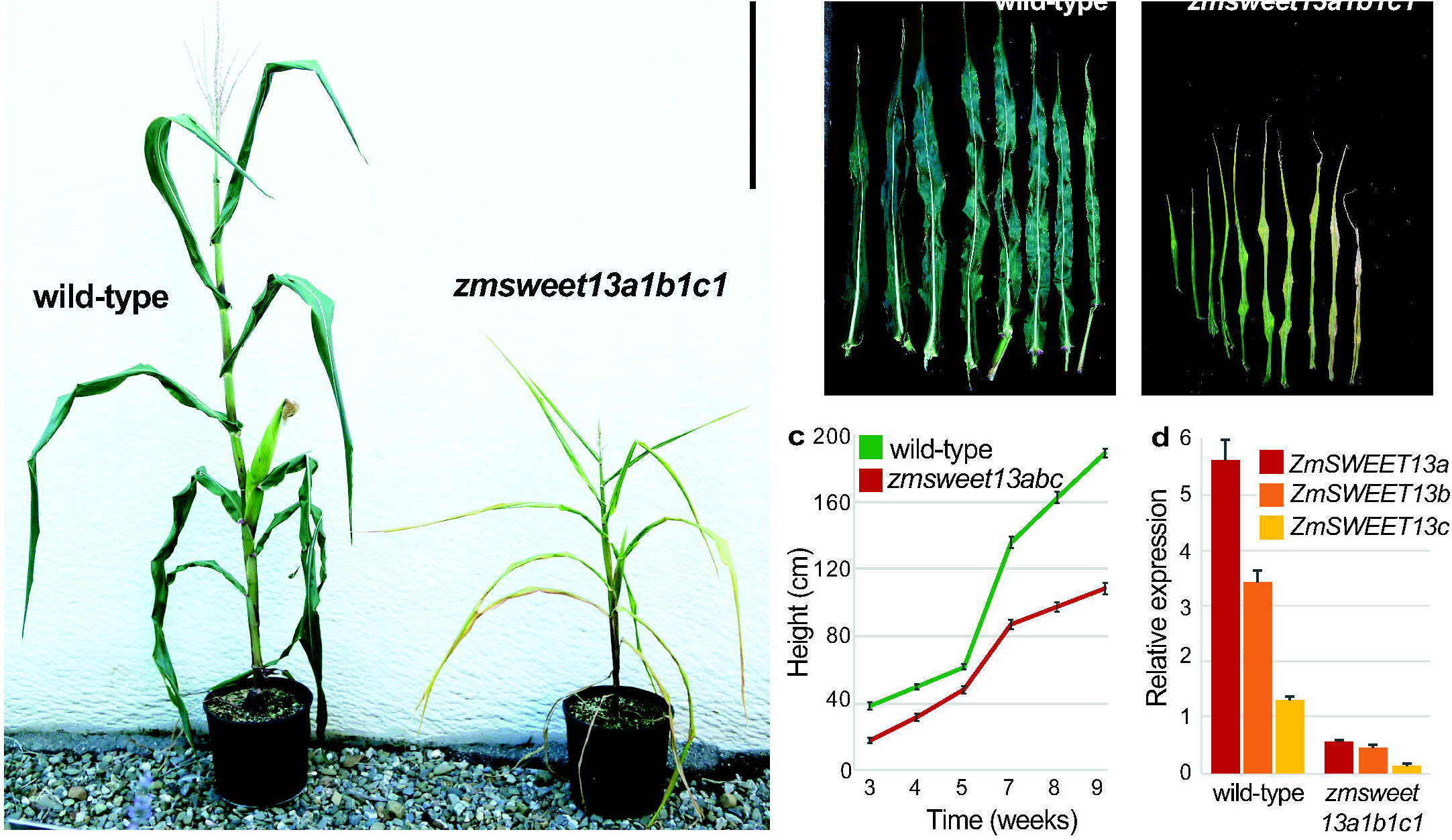
*ZmSWEET13* paralogs triple mutant characterization. **(A)** Photograph of fully mature wild-type and same-age *zmsweet13a1b1c1* triple mutant, showing reduced growth and leaf chlorosis. Bar: 50 cm. **(B)** Leaves comparison of plants presented in Fig. 2a, showing *zmsweet13a1b1c1* reduced growth and chlorosis. **(C)** Height quantification throughout wild-type and triple mutant life cycle in the greenhouse (mean ± s.e.m., n=17 and 15). **(D)** Relative expression (by qRT-PCR) of *ZmSWEET13* paralogs in maize flag leaf from wild-type and *zmsweet13abc*. Samples were harvested at 4:00 pm (mean ± s.e.m., n=3 technical replicates with expression normalized to 18S levels, repeated independently five times with comparable results).

**Figure 3.**
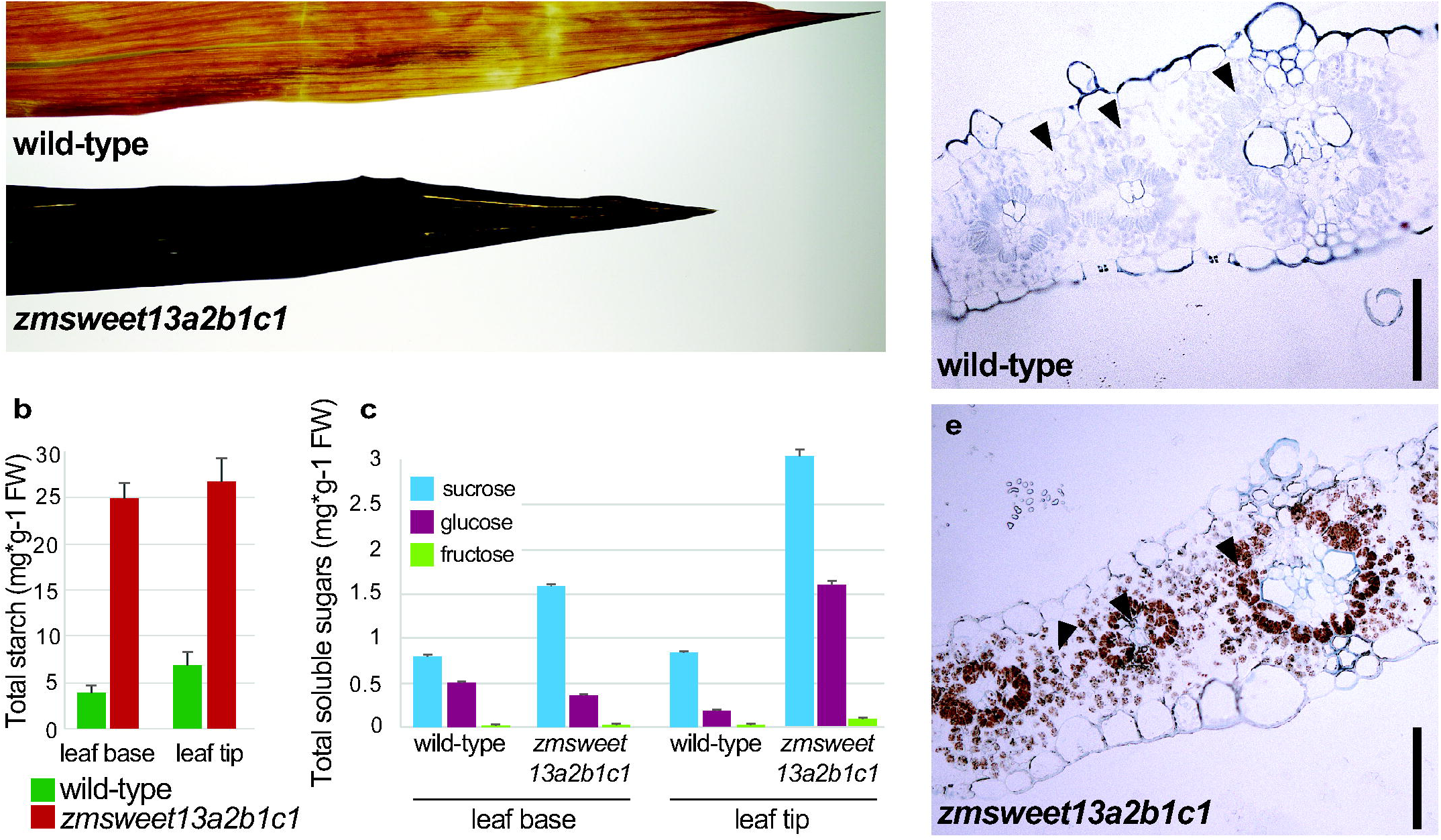
*zmsweet13abc* starch and soluble sugar accumulation. **(A)** Flag leaves were collected at dawn (7:00am), cleared with boiling ethanol and stained for 15 min with a saturated IKI (Lugol’s solution). **(B)** Starch quantification of leaves displayed in (A). The triple mutant shows 5x more starch in the leaf tip and blade, while no significant differences where measured in the sheath (mean ± s.e.m., n=3 technical replicates, repeated independently three times with comparable results). **(C)** Soluble sugar measurements of wild-type and *zmsweet13a2b1c1* in maize flag leaf sections. The triple mutant shows higher sugar concentration in both the tip and the base of the leaf. Samples were harvested at 7:00 am (mean ± s.e.m., n=3 technical replicates, repeated independently three times with comparable results). **(D)** Cross-sections of wild-type and triple mutant leaves harvested at the end of the night, fixed, embedded and stained with Lugol’s solution for starch. Mutant *zmsweet13a2b1c1* accumulates high amounts of starch in both mesophyll and bundle sheath cells. Bar: 75 mm.

Despite the severe defects, triple mutant plants grown in the greenhouse (as well as a subset in the field) exported sufficient sugars from leaves to produce viable seeds. A possible explanation for the viability of the triple mutants could be compensation by other sucrose-transporting clade III SWEETs. To test this hypothesis and to obtain insights about possible physiological changes in the mutants, we performed an RNA-seq on flag leaves of wild-type (Hi-II outcrossed once to B73) and triple mutant plants (Hi-II background). Notably, we did not observe significant enrichment of mRNA of clade III *SWEETs*, arguing against transcriptional compensation by other clade III *SWEETs* (**Fig. S9**). Consistent with impaired photosynthesis and chlorosis, mRNA levels of multiple genes encoding functions in the light harvesting complex and chlorophyll/tetrapyrrole biosynthesis were substantially reduced in triple mutants (**Fig. S10** and **Fig. S11**). Furthermore, in line with the accumulation of starch and soluble sugars in leaves, we found that transcripts related to carbohydrate synthesis and degradation were affected in the triple mutants (**Fig. S12**).

A recent study has found that the *Arabidopsis* homolog AtSWEET13 (although phylogenetically not the closest homolog of ZmSWEET13) can also transport the plant hormone gibberellin (Kanno *et al.*, 2016). The observed phenotypes for the triple zm*sweet13 knock out* mutants in maize are consistent with a primary role in sucrose transport and distinct from the ones observed in the *Arabidopsis sweet13;14* double mutant; i.e., male sterility, increased seedling and seed size (Kanno *et al.*, 2016).

To determine if variation in the *ZmSWEET13* genes may account for differences in agronomically important traits in existing maize lines, we conducted a genome-wide association study (GWAS) using phenotypic traits obtained from maize diversity panel (Flint-Garcia *et al.*, 2005). We obtained genotypic data from maize HapMap3 SNPs (Bukowski *et al.*, 2015) and filtered out SNPs having minor allele frequency < 0.1 and missing rate > 0.5, leaving ∼13 million SNPs for analyses. We performed GWAS using a mixed linear model approach (Zhou & Stephens, 2012), where kinship calculated from the genome-wide SNPs was fitted as the random effects. The SNPs that passed the FDR threshold of 0.05 and showed linkage disequilibrium (R^2^ > 0.8) with *ZmSWEET13s* genes were considered significant associations. SNPs in *ZmSWEET13s* were significantly associated with ear-related traits (i.e. ear rank number and ear height) and developmental traits (i.e. days to silk, days to tassel, middle leaf angle, and germination count) (**Fig. S13** and **Fig. S14**). While these results are compatible with a key role of *ZmSWEET13s* in carbon allocation, it will be necessary to determine whether polymorphisms in these genes or flanking regions are causative for these traits.

## Discussion

Phloem sap of many monocots and dicots contains very high sucrose concentrations, and it is thought that this gradient creates the drive for phloem translocation of sucrose and all other molecules in the phloem sap. Inhibition of the production of the SUT1 protein, by RNAi or mutations, typically leads to stunted growth and accumulation of carbohydrates in leaves (Riesmeier *et al.*, 1994; Bürkle *et al.*, 1998). Chlorosis and inhibition of photosynthesis, which often accompany the general growth defects, may either be due to feedback inhibition of photosynthesis or may be a consequence of nutrient deficiencies caused by the reduced supply of carbohydrates to the root system (Ainsworth & Bush, 2011). SUTs function as sucrose/H+ symporters and appear to fulfill two roles: (i) loading of the SECC with sucrose in source leaves, and (ii) retrieval of sucrose that diffuses out of the SECC as a consequence of the high sucrose concentration in the SECC relative to surrounding tissues. SUTs import sucrose from the cell wall space, implying the existence of transporters that efflux sucrose into the cell wall space preceding uptake by SUTs. AtSWEET11 and 12 are candidates for this role in *Arabidopsis*: they appear to function as uniporters and can thus serve as cellular efflux systems. Both SWEETs were highly expressed in leaves, localize to the phloem parenchyma, and *atsweet11;12* mutants were smaller and accumulated starch in leaves. However, the phenotype was relatively weak, implying leaky mutations, compensation by other transporters, or the coexistence of other phloem loading mechanisms. Other mechanisms could include symplasmic transport, or yet unknown processes. Lastly, is possible that SUTs play a major role in sucrose retrieval along the path in addition to their role in phloem loading, while ZmSWEET13’s are thought to be only involved in sucrose efflux prior to SUT1 uptake into phloem. Would a mutant of *ZmSWEET13* present weak defects similar to those in *atsweet11;12* plants, or a more severe phenotype equivalent that in *zmsut1* mutants?

Here, we show that maize has three paralogs in clade III of the SWEET family that are among the most highly expressed genes in leaf tissue. They derive from relatively recent gene duplication events: sorghum and wheat have three copies per genome, while Brachypodium and rice each have only one. The comparatively high number of *SWEET13s* had been attributed to specific roles in C4 photosynthesis, however the presence of three *SWEET13s* in wheat puts this interpretation into question. Evidence that the maize SWEET13s cooperate in phloem loading is based on two key observations: a severe growth defect similar to that of *zmsut1* mutants, and massive accumulation of free sugars and starch in leaves. These phenotypic effects are also similar to the RNAi phenotypes in potato and tobacco (Riesmeier *et al.*, 1994; Bürkle *et al.*, 1998). The observed growth defect in maize is much more severe than that of the *atsweet11;12* mutant in *Arabidopsis,* and comparable to that of the *zmsut1* mutant (Slewinski *et al.*, 2009; Chen *et al.*, 2012). We thus propose that the three ZmSWEET13s and ZmSUT1 play dominant roles in phloem loading.

It is noteworthy, that the combined *zmsweet13abc* mutations are not lethal, since the plants still produce fertile viable offspring, implying additional mechanisms for phloem loading in maize. While it is possible that other transporters might compensate, it is unlikely that other SWEETs take over such roles, as judged by the lack of induction of other clade III SWEET genes in the mutants. Therefore, maize likely uses either symplasmic or other loading mechanisms in parallel.

It is still not clear whether *SWEET13* triplication mainly serves to increase the amount of SWEET proteins in the same cells (e.g. phloem parenchyma), or if each SWEET13 transporter mediates efflux from a specific cell type and loading is achieved in a multi-tier fashion. This question is of particular interest because *in situ* hybridization experiments identified SUT1 in companion cells, xylem and phloem parenchyma, as well as the bundle sheath. To address this question, we generated translational reporter gene fusions and included the first three introns. However GUS activity or GFP fluorescence were not detectable in any of the transformants carrying fusions for any of the three SWEET13s (data not shown). We therefore hypothesize that additional elements are required for proper expression of these constructs.

Another interesting question is whether maize can serve as a model for phloem loading in rice, barley and wheat. Surprisingly, RNAi of the rice homolog of *ZmSUT1* did not lead to a detectable effect on the phenotype of the sporophyte (Ishimaru *et al.*, 2001). Thus, it remains a matter of debate whether rice uses predominantly apoplasmic and symplasmic or other mechanisms simultaneously. It will therefore be important to study the role of SWEET homologs in rice and other crops. It is noteworthy in this context that OsSWEET11 and 15 are expressed preferentially in the caryopsis and act as key players in apoplasmic unloading processes in developing rice grains (Yang, parallel submission)(Ma *et al.*, 2017).

Finally and most importantly, it will be interesting to see whether overexpression of clade III SWEETs in leaves of cereals may help to increase yield potential.

## Acknowledgments

We thank Joelle Sasse, Bi-Huei Hou, Aurélie Grimault, Franklin Talavera-Rauh, Xiao-Qing Xu, Grayson Badgley and Guido Grossmann for experimental support during various analyses. We thank Ari Kornfeld (Dep. Global Ecology, Carnegie Science) for help and advice in photosynthesis measurements. This work was made possible by research grants from Syngenta Crop Protection, LLC (WBF), the Office of Basic Energy Sciences of the US Department of Energy (DE-FG02-04ER15542; WBF), and the National Science Foundation (IOS-1258018; BY, WBF). WF’s lab is supported by the Alexander von Humboldt Foundation.

## Author Contributions

MB, SNC, BY, WBF and DS conceived and designed experiments. MB, MH, TH, SNC, DL, BL and DS performed experiments. MB, TH, BY, WBF and DS analyzed the data. DS and WBF wrote the manuscript, MB and BY helped with revisions.

## Competing Financial Interests

The authors (DS and WBF, on behalf of Carnegie Science) have filed a PCT patent application based in part on this work with the US Patent and Trademark Office.

